# Structural-functional connectivity mapping of the insular cortex: A combined data-driven and meta-analytic topic mapping

**DOI:** 10.1101/2021.07.07.451405

**Authors:** Benjamin Klugah-Brown, Pan Wang, Yuan Jiang, Benjamin Becker, Peng Hu, Lucina Q. Uddin, Bharat Biswal

**Affiliations:** The Clinical Hospital of Chengdu Brain Science Institute, MOE Key Laboratory for Neuroinformation, Center for Information in Medicine, School of Life Science and Technology, University of Electronic Science and Technology of China, No.2006, Xiyuan Ave, West Hi-Tech Zone, 611731, Chengdu, China; Department of Biomedical Engineering, New Jersey Institute of Technology, Newark, NJ 07102, USA; Department of Psychiatry and Biobehavioral Sciences, University of California Los Angeles 760 Westwood Plaza Los Angeles, CA 90095

**Keywords:** DTI, Human Connectome Project, Insular, Resting-state functional connectivity

## Abstract

In this study, we examined structural and functional profiles of the insular cortex and mapped associations with well-described functional networks (FNs) throughout the brain using diffusion tensor imaging (DTI) and resting-state functional connectivity (RSFC) data. We used a data-driven method to independently estimate the structural-functional connectivity of the insular cortex. Data were obtained from the Human Connectome Project comprising 108 adult participants. Overall, we observed moderate to high associations between the structural and functional mapping scores of three different insular subregions: the posterior insula (associated with the sensorimotor network: RSFC, DTI = 50% and 72%, respectively), dorsal anterior insula (associated with ventral attention: RSFC, DTI = 83% and 83%, respectively), and ventral anterior insula (associated with the frontoparietal: RSFC, DTI = 42% and 89%, respectively). Further analyses utilized meta-analytic decoding maps to demonstrate specific cognitive and affective as well as gene expression profiles of the three subregions reflecting the core properties of the insular cortex. In summary, given the central role of the insular in the human brain, our results revealing correspondence between DTI and RSFC mappings provide a complementary approach and insight for clinical researchers to identify dysfunctional brain organization in various neurological disorders associated with insular pathology.

## Introduction

Determining the intrinsic structural and functional interplay of brain regions critical for cognition is vital for understanding typical and atypical human behavior. Capitalizing on the development of functional magnetic resonance imaging (fMRI) and diffusion tensor imaging (DTI) (P. Wang et al., 2021) enables examination of the functional interactions of discrete brain regions and their structural underpinnings at the systems level. Although these two imaging modalities have been combined for exploring the characteristics of the brain independently [see review in (Rykhlevskaia, Gratton, & Fabiani, 2008)], most studies focus on measuring brain activity in response to a specific task, where interpretations are often limited to the brain systems engaged by the specific experiment. To study the global and general characterization of the associations between the two modalities, an intrinsic whole-brain approach might be more appropriate. Consequently, DTI and resting-state functional connectivity (RSFC) are increasingly used in a complementary fashion to task activation-based approaches (Baird, Colvin, VanHorn, Inati, & Gazzaniga, 2005; Kelly et al., 2012; Toosy et al., 2004). Motivated by recent studies (Long et al., 2017; P. Wang et al., 2021), the present study explored the correspondence between the structural and functional connections of the human insular cortex and associated mappings to well-described intrinsic functional networks (FNs) of the brain.

The insula is an important functional and anatomical integration hub, with connections to multiple cortical and subcortical regions. Numerous previous studies have explored functional and structural connectivity profiles of the insula independently(Cauda et al., 2011; Cauda & Vercelli, 2013; Cerliani et al., 2012; Deen, Pitskel, & Pelphrey, 2011; Kurth, Eickhoff, et al., 2010; Jason S. Nomi et al., 2016). With reference to white matter tracts, several cortical and subcortical regions, including the frontal, parietal, hippocampus, and amygdala, were shown to be structurally connected to the insula (Showers & Lauer, 1961). Furthermore, using diffusion-weighted imaging, human studies have shown similar connectivity between the posterior/anterior insula (PI/AI) and the anterior cingulate, frontal, parietal, and sensorimotor regions (Cloutman, Binney, Drakesmith, Parker, & Lambon Ralph, 2012; Ghaziri et al., 2017). With respect to multimodal investigations, an early study by Kelly and colleagues explored the insular cortex using three independent imaging modalities: task-evoked coactivation, intrinsic functional connectivity, and gray matter covariance (Kelly et al., 2012). They showed that the underlying brain organization consistently reflected the structural-functional tripartite cytoarchitecture of the insula.

In the present study, we examined the insular-cortical systems by capitalizing on two imaging modalities (resting-state fMRI and DTI) in healthy adults. Though several independent structural and functional connectivity studies of the insula have been conducted, previous studies have not addressed the correspondence of structural connectivity (DTI) and RSFC of the insular cortex and associated mapping to intrinsic FNs, thereby limiting overall understanding of the complementary association between the fiber tractography connectivity and the resting-state imaging methods and how they may enhance clinical research in various mental disorders. Therefore, mapping the structural and functional connectivity overlaps of insular subregions to the large-scale intrinsic networks of the whole brain could enable the description of well-defined characteristics underlying the varied functions of this multipurpose cortical region.

In this work, we explored these associations using a data-driven method to first test the uniformity in the human insular cortex and its mapping to well-described intrinsic FNs. We also used meta-analytic topic mapping to examine the cognitive and behavioral domains associated with each insular subregion. The general significance of this present study is that, although studies have investigated the individual imaging modalities with respect to the insula, our approach shows novelty in two aspects; firstly, in relation to the insula, we answer the question regarding the extent to which the structural profiles derived from DTI compares with functional properties obtained from RSFC. Notwithstanding the specific assumptions underlying each of the two imaging techniques, our independent structural and functional connectivity parcellation which demonstrated good overall concordance provides more insight into unique subregional differentiations relevant for research and intervention in insula-related disorders (Naqvi & Bechara, 2009; Wiebking et al., 2015). Secondly, in previous meta-analytic studies (Cauda et al., 2012; Chang, Yarkoni, Khaw, & Sanfey, 2013; Uddin, Kinnison, Pessoa, & Anderson, 2014), authors did not consider the correspondence of structural and functional insular subregions which could underpin the assumption that synaptic connections collaborated to provide a functional role in the FNs. In addition, while it’s important to track the neural and physical boundaries of the insula, the cognitive/behavioral and gene information have been not been fully updated, we, therefore, highlight these properties in this paper. Thus, we first identified these overlapped subregions and then proceeded to decode their subsequent coactivations. In all, the study not only reinforces our understanding of the insula but also provides a novel approach for investigating structural-functional connectivity profiles of other brain regions.

## Materials and Methods

### Data Acquisition

We obtained 119 healthy participants from the HCP dataset, of which 108 (11 DTI participants’ datasets were of different parameters) were used for the entire analysis, similar to our previous study (P. Wang et al., 2021). Although the dataset included resting-state fMRI with four runs (rfMRI_rest1_LR, rfMRI_rest1_RL, rfMRI_rest2_LR, rfMRI_rest2_RL), for T1-weighted images, we used a single run (rfMRI_rest1_LR) similar to the above-mentioned previous study. See Supplementary Materials and Methods for acquisition details.

### Functional and DTI Data Preprocessing

The functional images were preprocessed using a combination of toolboxes: Data Processing Assistant for Resting-State fMRI (http://rfmri.org/DPARSF) and SPM12 (http://www.fil.ion.ucl.ac.uk/spm/software/spm12). The DTI preprocessing involved: normalization of b0 image intensity with the following correction; EPI distortions, eddy current distortions, participant movements, and gradient nonlinearities (Glasser et al., 2013). All preprocessing steps were similar to our previous work (P. Wang et al., 2021). See Supplementary Materials and Methods for preprocessing details.

### Identification of Insula Functional Subregional Networks

We obtained seven functional cortical networks from Yeo’s template (Yeo et al., 2011) (downloaded from https://surfer.nmr.mgh.harvard.edu/fswiki/CorticalParcellation_Yeo2011): VSN, SMN, DAN, VAN, LIMB, FPN, and DMN. The insula mask was obtained from Harvard-Oxford cortical and subcortical structural atlases (https://neurovault.org/collections/262/). We excluded the insula voxels within the Yeo networks. Then, for each participant, the partial correlation coefficient was computed between every voxel time series within the insula mask and the averaged time series of each cortex network, while controlling the impact of the temporal contributions from the remaining six cortex networks. Fisher’s Z transform was performed on all correlation coefficients. To obtain the associated subregions, we projected the correlation coefficient to each voxel corresponding to the insula. A statistical t-map of connectivity pattern was obtained by calculating a one-sample t-test across all participants. To obtain the insular subregions corresponding to each intrinsic FN (seven FNs), we used the winner-take-all algorithm by selecting the most similar profile of connectivity for each insular voxel. As the winner-take-all method assumes that one voxel within the insula only corresponds to a single network, we performed a statistical correction of the one-sample t-test map of each cortex network as a region of interest (ROI) (P < 0.05, familywise error [FWE]-corrected) to obtain a more comprehensive understanding of the relation between the insula and FNs.

### Identification of Insula Structural Subregional Networks

We selected the seven Yeo intrinsic FNs and insula mask as ROIs for the DTI analysis. Before tracking the fibers connecting the FNs and insula, these FNs and insula mask were resampled to standard MNI diffusion space with a voxel size of 1.25 × 1.25 × 1.25 mm3. For each participant, we extracted the fiber bundles from the whole-brain fibers connecting each FN and insula, which was implemented in TrackVis (www.trackvis.org). Subsequently, we accumulated these fiber bundles corresponding to each FN across all participants to establish a population-based probabilistic map of commissural tracts using the approach of Park et al. (Park et al., 2008). The computations resulted in population probabilistic maps whose connectivity value corresponded to the number of samples with a trajectory from the insula mask through voxels and then to the cortical target mask. Essentially, a voxel was identified as part of a tract if it belonged to at least 15% (Bennett, Madden, Vaidya, Howard, & Howard, 2011; Khalsa, Mayhew, Chechlacz, Bagary, & Bagshaw, 2014) of the participants. The procedure was aimed at validating the harmony between the anatomical properties and the functional connectivity.

### Examining the Significant Overlap Between DTI and RSFC Mappings

Using the Monte Carlo method (D. Zhang, Snyder, Shimony, Fox, & Raichle, 2010), we examined the significant overlap of insular subregions between the DTI and RSFC mapping. To ascertain these overlaps, we used all insular voxels and assigned each insular voxel a binary value (0 and 1). We then determined the proportion of voxels with a value of 1 by computing a 15% threshold of the total NOV in the DTI and RSFC (statistically significant threshold at P < 0.05, FWE-corrected). Overlap between the DTI and RSFC mapping was defined as the number of simulated voxels containing a value of 1 in both maps. A random permutation was performed 10,000 times, which assigned a value of 1 to the DTI and RSFC voxels. A statistically significant level was calculated by estimating the null distribution and NOV in the overlap. We categorized the overlaps using the following criteria: low (0–0.19), low–moderate (0.2–0.39), moderate (0.4–0.59), moderate-high (0.6–0.79), or high (0.8–1) as defined in (2).

### Meta-Analytic Topic Mapping and gene expression

We capitalized on meta-analytic topic mapping insular subregions to facilitate a functional characterization and differentiation by means of the Neurosynth database (https://neurosynth.org/; (Yarkoni, Poldrack, Nichols, Van Essen, & Wager, 2011). To further determine separable biological profiles we also examine the gene expression related to the specific subregions identified. Here, we first defined the center of mass for each subregion on either side of the hemisphere, and next created unthresholded maps comprising two different statistical inference maps: the uniformity test map and the association test map (false discovery rate [FDR] = 0.01). The uniformity map indicates the extent to which each brain voxel is consistently activated in studies for a specific term. The association test map indicates whether the activity in a voxel occurs more specifically for studies using the term of interest than for studies not using this term. Using these maps, a decoding function was applied which generates terms associated with the subregion, next we extracted the top ten (10) terms by means of their correlation values. In addition, we employed the BAT software (Liu et al., 2019) to identify the associated genetic expression for the left and right insular. The top ten (10) gene symbols were extracted and for each symbol, we extracted the correlation values associated with the subregions indicating the extent to which subregional terms associate with genetic information.

## Results

### Identification of Functional Insular Subregional Networks

Seven cortical intrinsic FNs were defined based on Yeo’s template (Yeo et al., 2011): visual network (VSN), sensorimotor network (SMN), dorsal attention network (DAN), ventral attention network (VAN), limbic network (LIMB), frontoparietal network (FPN), and default mode network (DMN) (Figure. 1A). Figure 1B depicts the distinct insular subregions with maximum voxels functionally connected to the intrinsic FNs and computed through the winner-take-all algorithm. The algorithm takes into account one voxel within the insula per network. As a result, only six distinct insular subregions were connected to the six networks, leaving out the VSN (Figure 1B, row 2, column 1). For low-order intrinsic FNs, we found that the SMN connected with the left/right/bilateral PI (left, dorsal granular; right, hypergranular; Figure 1B, red). The DAN was also connected to the left PI (dorsal granular, Figure 1B; blue). The subregion corresponding to the VAN and LIMB showed connectivity distribution in the bilateral dAI (dorsal granular and ventral dysgranular, Figure 1B; yellow and cyan, respectively). Maximal connectivity with the FPN and DMN was mostly found in the left/right vAI (left; ventral granular, Figure 1B, orange) and right vAI (ventral granular, Figure 1B; green), respectively.

**Figure 1.**
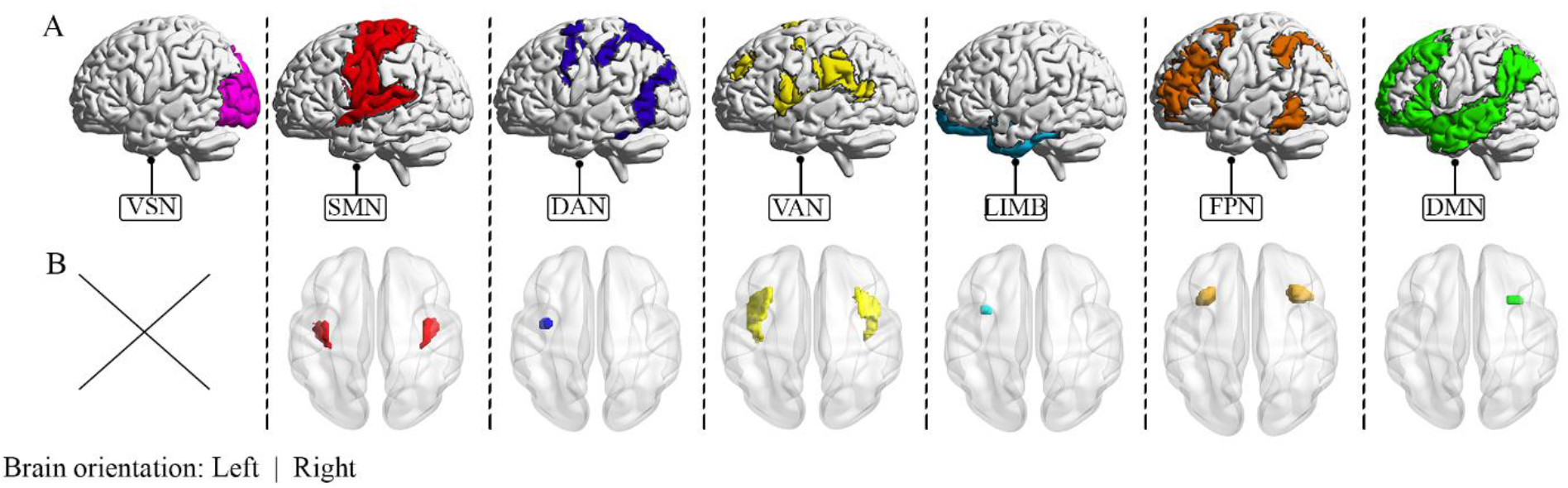
Winner-take-all projections of the subregional insula maps. (A) The cortex is partitioned based on major anatomical landmarks into seven nonoverlapping FNs. (B) Each insular subregional voxel is labeled with the color of the FN that produced the highest partial correlation values using RSFC. The empty cell indicates no voxel per the criterion of the winner-take-all algorithm; the subregion had too few voxels to survive the iteration.

Figure 2 demonstrates the computed on the voxels-wise structural and functional subregions. For functional connectivity, we observed a distinctive pattern for both the “winner-take-all” and the FWE. There were six (6) subregions each corresponding to the networks except for the visual network due to a very low number of voxels, as shown in Figure 2 first row (indicated with “none”) identified with the FWE. Specifically, Maximal connectivity was observed in vAI subregions (vAI; left NOV = 206, right NOV = 222; vAI, left NOV = 33, right NOV = 21) connected to the DMN and FPN, respectively (Figure 2, rows 6, 7). Two dAI (left NOV = 990, right NOV = 968;, left NOV = 207, right NOV = 148) were maximally connected to the VAN and LIMB, respectively (Figure 2, rows 4 and 5). For the PI, we found two subregions (left NOV = 419, right NOV = 496; left NOV = 48, right NOV = 48). Table S1 shows the detailed peak coordinate of each insular subregion.

**Figure 2.**
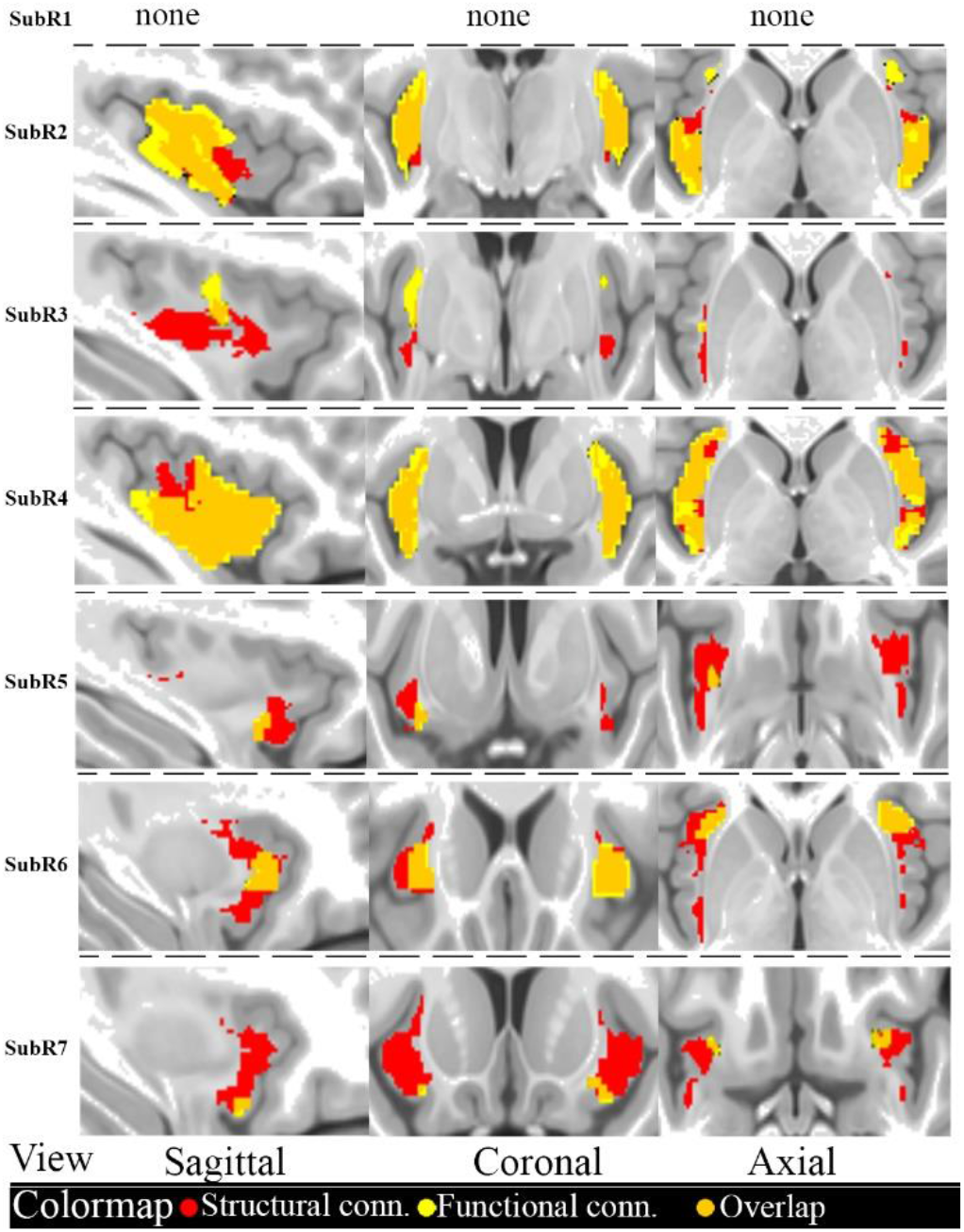
Functional and structural connectivity of insular cortex. Structural connectivity derived from probabilistic tractography demonstrated specificity of tracking white matter fiber tracts between the insula and each FN; all structural results were obtained after thresholding to 15% of the total participant voxels (color-coded; red). Functional connectivity profile; each FN (color-coded; yellow) shows specific correlations with distinct insular subregions. The results were thresholded using FWE following the winner-take-all algorithm shown in Figure 1B. Structural and functional mapping demonstrated significant overlap in their connectivity profiles (color-coded; orange). SubR; subregion

### Structural Connectivity Between the FNs and Insular Subregions

For each participant, fiber bundles connecting each of Yeo’s seven cortical networks and insular regions were extracted using TrackVis (www.trackvis.org). Through a 15% threshold of the participants’ voxel allocation, we identified significant mapping between the insular subregion and the FNs similar to the functional connectivity. Figure 2 (red color map) demonstrates the DTI connectivity of each subregion (row 6, left NOV = 1437, right NOV = 833; row 7, left NOV = 1686, right NOV = 832) categorized as vAI and connected to the VSN, FPN, and DMN, respectively. Also, the dAI subregions (row 4, left NOV = 3196, right NOV = 2339; row 5, left NOV = 733, right NOV = 319) were maximally connected to the VAN and LIMB, respectively. For PI, two subregions (row 2, left NOV = 1882, right NOV = 1542; row 3, left NOV = 237, right NOV = 97) demonstrated connectivity with the SMN and DAN. Details of the peak insula coordinates are shown in Table S2.

### Structural-functional Connectivity Overlap Between DTI and RSFC

The differences in the underlying physical characteristics of the two methods notwithstanding, there were significant (P < 0.0001) overlaps in the connectivity profile of each subregion mapping to each intrinsic FN. Figure 2(orange color map) shows the overlap between the RSFC and DTI results (Table S3 shows the detailed peak coordinates for the identified subregions). The insula connectivity demonstrated a similar pattern of mapping to the FNs. The DTI and RSFC significantly overlapped largely with all regions and mapped to all FNs except the VSN. The overlap of the insular subregion corresponding to the DAN between the DTI and RSFC was exceptionally low (<7%, P = 0002) for both the winner-take-all algorithm and the FWE computation. Based on the following criteria: low (0–0.19), low–moderate (0.2–0.39), moderate (0.4–0.59), moderate-high (0.6–0.79), or high (0.8–1) as defined in (2), three significant insular regions showed greater than moderate overlap (Figure 2, row 2, 4, 6,, Table 1), suggesting the largest overlap between DTI and RSFC (PI-SMN [PI related to the SMN], RSFC = 50%, DTI = 72%; dAI-VAN [dAI associated with the VAN], RSFC = 83%, DTI = 83%; vAI-FPN [vAI associated with the FPN], RSFC = 42%, DTI = 89%).

**Table 1.** For each functional map, percent overlap was calculated as the number of voxels that overlap between DTI and RSFC divides by the total number of voxels that exceed the threshold, respectively. The significance of overlap (p-value) between DTI and RSFC is obtained by using the Monte Carlo method. Bold values represent larger overlaps. All values are converted into percentages

|  | Winner takes all |  | FWE |  | significance |
| --- | --- | --- | --- | --- | --- |
| FNs | RSFC overlap (%) | DTI overlap (%) | RSFC overlap (%) | DTI overlap (%) | (P -value) |
| VSN | 0 | NaN | 0 | 0 | NAN |
| SMN | **40** | **71** | **50** | **72** | **P<0.0001** |
| DAN | 0.7 | 5 | 2 | 5 | P=0.002 |
| VAN | **70** | **87** | **83** | **83** | **P<0.0001** |
| LIMB | 2.5 | 70 | 12 | 38 | P<0.0001 |
| FPN | **2.5** | **96** | **42** | **89** | **P<0.0001** |
| DMN | 23 | 100 | 2.3 | 95 | P<0.0001 |

Based on the percentage overlaps in, Table 1, we obtained the overlap maps computed through the winner-take-all algorithm and the FWE for the three distinct insular subregions, namely the PI (Brodmann area 22), dAI (Brodmann area 44), and vAI (Brodmann area 4) (Figure 3A, B, C); the cytoarchitecture of the three regions (Figure 3D) resembled previously reported cluster analysis results (Deen et al., 2011).

**Figure 3.**
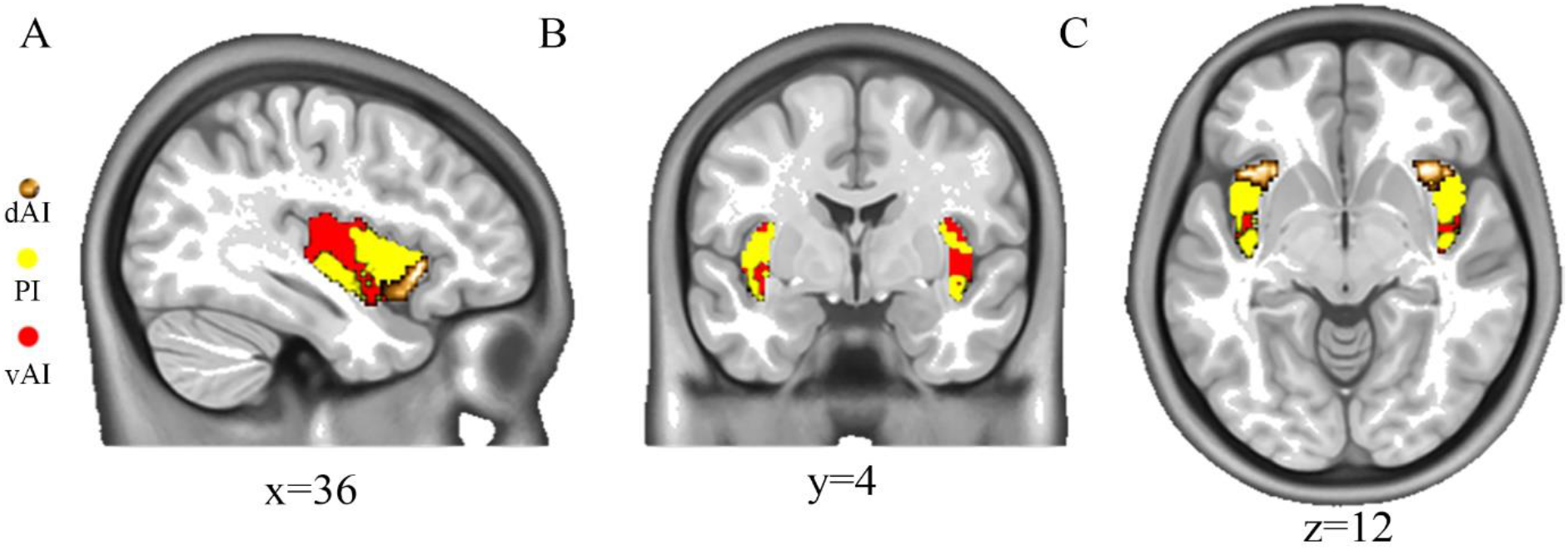
Distinct insular subregions per significant overlap. (A–C) Winner-take-all computed for three distinctive insular subregions; each color signifies a specific voxel. Each voxel was mapped to the FNs as follows: PI-SMN, dAI-VAN, and vAI-FPN. Results displayed in the orthogonal view

### Topic Mapping of Insular Subregions

We examined the relationships between the insular subregions (left and right) and top 10 cognitive and behavioral terms as well as their related genetic expressions. Figure 4 demonstrates that left and right insular subregions have both distinct and common associative terms. Generally, “Pain” is associated highly with dAI and PI, while “working memory or memory” is associated highly with vAI. Moreover, top terms on the left for each subregion demonstrated higher correlations relative to the right. In addition, five gene symbols (THEMIS, SAMD3, KRT17, LINC00238, KRT16P2, and DAPP1) were expressed in both left and right insula, but with different levels of associations for each hemisphere of the subregions. In general, compared to dAI and PI for both left and right, the vAI expressed a lower association in all the gene symbols (Figure A, B; third column). Across the subregions for both left and right insular, different symbols correlated highly with the region; for left dAI (LINC00238; r=0.084), left PI (DAPP1; r=0.198), and vAI (SLC26A4, r=0.018), for right dAI (KRT1; r=0.108), right PI (DAPP1; r=0.198) and right vAI (KRT1; r=0.03), all correlation were threshold as p=0.01.

**Figure 4.**
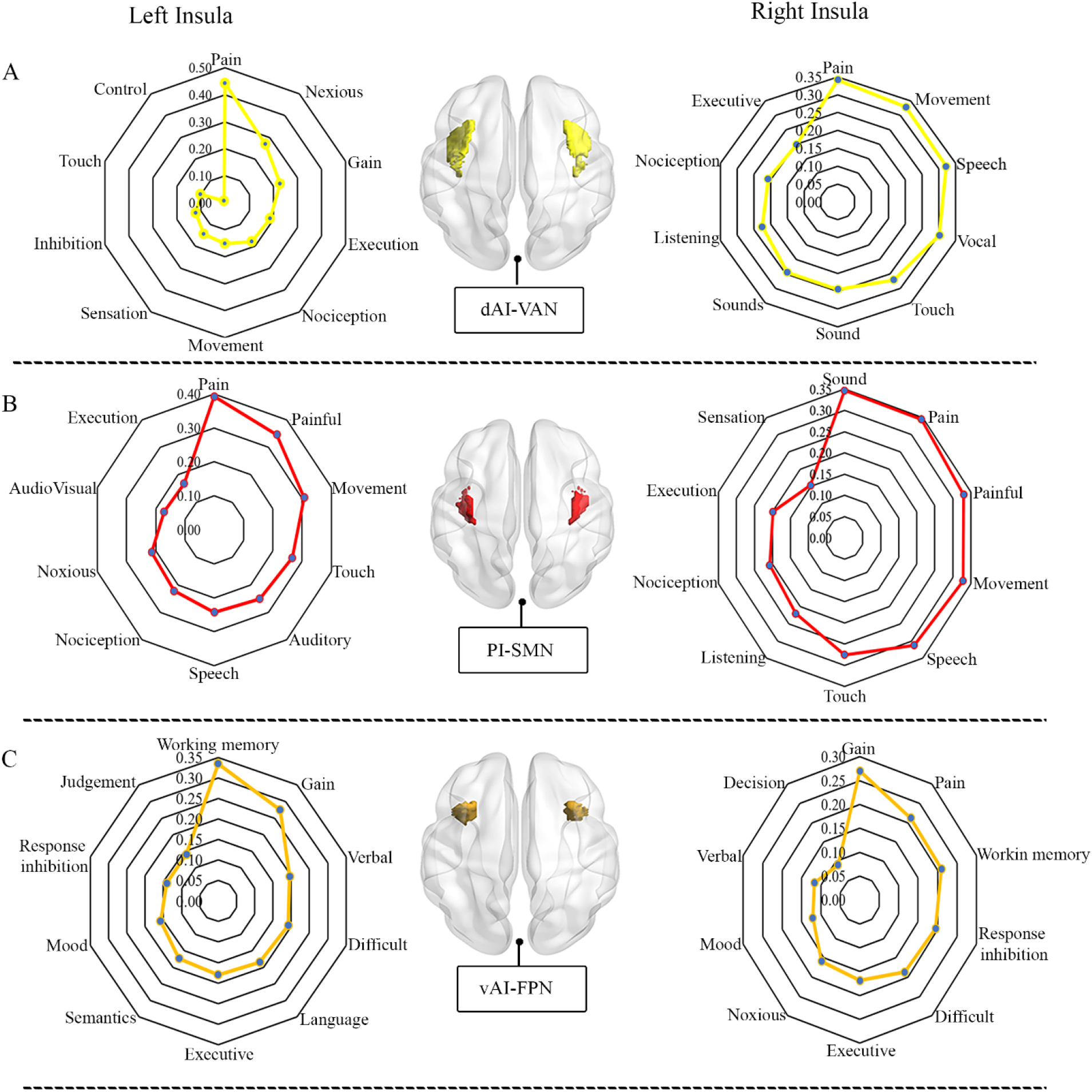
Meta-analytic topic mapping. Each term signifies cognitive and behavioral domains of studies in the database. Values displayed on the web represent the top ten (10) correlation of terms associated with subregions obtained in Figure 4. Significant association threshold at FDR = 0.01

**Figure 5.**
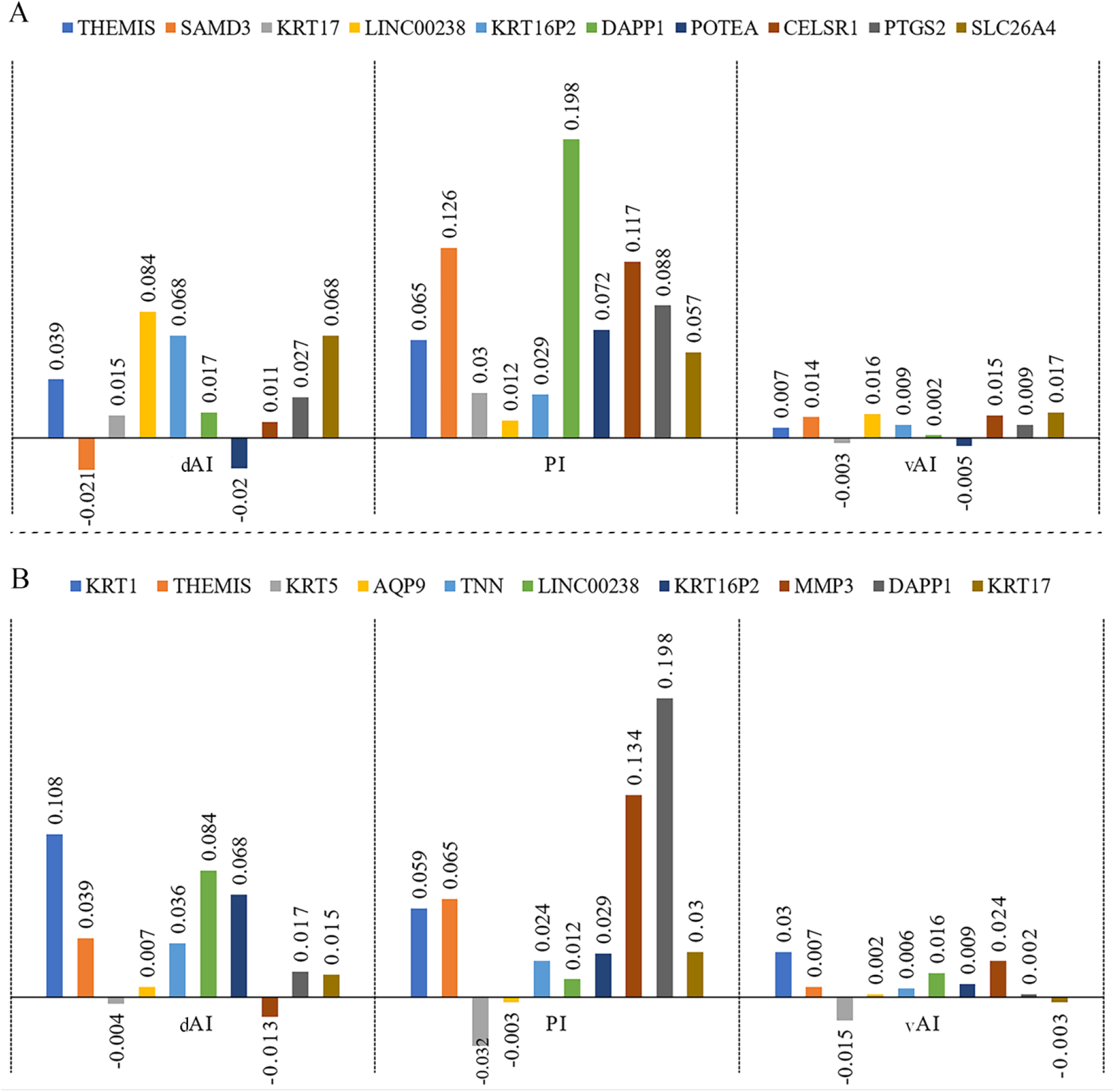
Genetic expression. (A and B) demonstrates the gene symbol expression associated with left and right insular subregion, respectively. Each bar shows the correlation values extracted for the top ten (10) symbols from the Neurosynth database. The significant threshold is set at p<0.01

## Discussion

In the present study, we demonstrated the correspondence between the structural and functional connectivity of insular subregions in the human brain and their mapping to well-defined intrinsic FNs. This approach highlights the basic structural-functional connectivity relationship of the tripartite architecture of the insula, with subregional distribution comprising the PI, dAI, and vAI. The generally moderate to high structural-functional correspondence revealed through the overlaps validated the potential mapping of insular subcortical connectivity via the two independent imaging methods and reflects the complementary relationship between the DTI and RSFC findings. Pairing the RSFC to the DTI tracts showed that the three distinctive insular regions were mapped significantly to three of the seven intrinsic FNs including SMN, VAN, and FPN, and indicating a direct physical connection. Notably, these regions are consistent with the cytoarchitecture of the insular cortex identified in earlier studies (Augustine, 1996; Chang et al., 2013; Kurth, Zilles, Fox, Laird, & Eickhoff, 2010; Mesulam & Mufson, 1982; Uddin, Nomi, Hébert-Seropian, Ghaziri, & Boucher, 2017) and linked to large-scale intrinsic FNs (Kurth, Zilles, et al., 2010; Yeo et al., 2011). Moreover, the meta-analytic topic mapping and the genetic expressions, performed in a relatively large database composed of terms and genetic symbol association, provides an additional perspective of the different insular subregions and how they are associated with various cognitive and behavioral domains across the networks as well as how they related to important genetic expressions in humans.

### Insular Connectivity Mapping to intrinsic FNs

Mapping the functional and structural correspondence of insular subregions is imperative for understanding the diverse functions of the insula and its relations with large-scale brain networks. There is a wide array of both structural (Menon et al., 2020; Shura, Hurley, & Taber, 2014) and functional (Jezzini, Caruana, Stoianov, Gallese, & Rizzolatti, 2012; J. S. Nomi, Schettini, Broce, Dick, & Uddin, 2018) profiles of the insula that have previously been published, providing insight into the connectivity of the distinct regions shown in the present study. The insular subregions revealed in the current work are consistent with that in previous functional (Nelson et al., 2010) and meta-analytical (Chang et al., 2013) studies. Regarding the correspondence of the two imaging modalities contributing to the connectivity of brain networks, Zhang and colleagues examined the differences and overlap between the structural and functional connectivity of the thalamus corresponding to cortical areas of the brain (Z. Zhang et al., 2011) and suggested a potential intersection of the thalamic system linking to specific cortical and subcortical regions across the brain. Jiang et al. showed a cortical connectivity profile that offset the dysfunctions associated with the motor perception networks of the gray matter in patients with schizophrenia (Jiang et al., 2019). In agreement with the abovementioned studies, our study demonstrates DTI intersecting with RSFC, suggesting a complementary relationship between each method.

In the present study, we observed three distinct structural and functional regions with moderate to high overlap that significantly mapped to specific FNs. Based on the structural and functional correspondence, we found different levels of percentage connectivity of overlap. For example, the VSN, DAN, and LIMB were highly significant (P < 0.0001) but showed lower overlap; in contrast, the SMN, VAN, and FPN showed significant overlap for both RSFC and DTI mapping (Table 1). We posit that the observed correlations of the RSFC and DTI fiber connection to the intrinsic FNs was due to the unique insular subregions mapping to the FNs and reflect the significant role the subregions play in cognitive functions. The greatest overlap using percentage voxel count was exhibited in mapping with the VAN (Figure 2, row 4, Table 1); the extent of the overlap suggested a strong complementary mechanism between the functional and structural connectivity with uniform correlation among the FNs and indicating that both methods contribute to overall characterization at the systems level.

Both the winner-take-all algorithm and the FWE demonstrated high structural connectivity mapping with the intrinsic FNs as compared with the RSFC, with the highest overlap occurring in the dAI mapping to the VAN. The variability in the level of correspondence between the two imaging modalities was due to DTI connectivity being fairly consistent over time (Z. Zhang et al., 2011). DTI represents physical connections or axonal tracts interconnecting distinct brain regions, with significant stability relative to functional connectivity, which in contrast represents correlated functional activity between brain regions and exhibits more variability over time (D. Zhang et al., 2010). Additionally, the two imaging methods were expected to yield different, yet complementary results due to their underlying characteristics. Interestingly, significant overlaps were found with a stricter threshold (P < 0.0001), suggesting that DTI and RSFC are comparable in terms of their abilities to detect meaningful connections between brain regions.

### Insular Subregional mapping to intrinsic FNs

Remarkably, the parcellation through the winner-take-all algorithm and the FWE demonstrated relatively robust clusters across the insular region. The distinct subregions were initially grouped in two major subregions: the AI and the PI (Figure 1, Table S1, S2). The overlap scores further grouped the region into three distinct subregions: PI, dAI, and vAI (Figure 3A-C, Table S3), consistent with that reported previously (Chang et al., 2013; Deen et al., 2011). Kurth and colleagues reported four functional divisions of the insula: PI (involved in sensorimotor processes), central part (involved in olfactory and gustatory processes), and the vAI–dAI (involved in cognitive and socioemotional processes) (Kurth, Zilles, et al., 2010); the functional distribution echoed that in the present study, with the central part further reorganized to the dAI (Figure 3B).

Functional and structural connectivity of the PI mapped significantly to the SMN (Figure 1A, B; column 2), with regional connectivity including primary/secondary motor and somatosensory similar to those reported in previous insular parcellation studies (Deen et al., 2011; Kelly et al., 2012). Interestingly, the observed structural-functional overlap reflected the widespread physical and neural complements implicated in the SMN. Prior functional studies using a data-driven approach reported that the regions found in the network showed precentral, postcentral, and sensorimotor areas (Beckmann, DeLuca, Devlin, & Smith, 2005), similarly in connectivity studies involving somatosensory and interoceptive processes (Craig, 2002; Kuehn, Mueller, Lohmann, & Schuetz-Bosbach, 2016; McGlone et al., 2002; Olausson et al., 2002; X. Wang et al., 2019). Combined, the abovementioned studies and our findings reflect the contribution to the primary regions involved in affective and somatosensory processes.

For the anterior insula, there were two further divisions: dAI and vAI, occupying the central and anterior sections of the insular cortex (Figure 3B, C), respectively. The dAI (Figure 3B) mapped to the VAN (Figure 1A, column 4), with high connectivity in the frontal, anterior cingulate, and parietal areas functionally consistent with the clustering method (Deen et al., 2011). The VAN is implicated in attentional and awareness stimuli (Indovina & MacAluso, 2007; Serences et al., 2005) during tasks. The structural-functional overlap (83% overlap, Figure 2; row 2 and 4, Table 1) demonstrated the consistency of the physical connectivity most relevant to the dAI. In addition, the dAI is functionally related to the brain areas involved in cognitive control processes (Dosenbach et al., 2007). In their clustering analysis, Deen et al. reported significant connectivity between the dAI and the regions involved in attentional control (Deen et al., 2011), which is in agreement with our results involving the VAN.

Ventrally in the AI, the results demonstrated significant (Figure 3C) mapping to the FPN (Figure. 1A, column 6), with significant overlap between DTI and RSFC, which includes the anterior cingulate cortex (ACC), posterior parietal cortex, and mid/superior frontal cortex. Unlike the dAI, the vAI exhibited a relative imbalance overlap, in which the maximum percentage was demonstrated in the DTI profile (Table 1); this may suggest the extent of the voxels from the neighboring dAI. Functionally, the FPN comprised mainly the ACC, frontal, and parts of the parietal regions. The significant correspondence between the structural-functional vAI mapping to the FPN suggests the role of salience detection and cognitive control (Seeley et al., 2007).

Additionally, Medford and Critchley reviewed the functional relationship between the AI and the ACC and highlighted the complementary relationships, representing sensory input and awareness processes (Medford & Critchley, 2010). The posterior part of the ACC has also been suggested to strongly correlate with the vAI, and exhibits salience characteristics (Marek & Dosenbach, 2018; Margulies et al., 2007; Seeley et al., 2007). Further, for emotions, pain from nociception is a vital sensory perception associated with both the vAI and dAI; the vAI is also associated with neuropathic pain (Ferraro et al., 2021). Although structural evidence through DTI is limited, its high overlap with RSFC suggested an important mapping between the vAI and the FPN.

### Meta-Analytic Topic Mapping and gene expression

The three subregions identified are consistent with previous studies (Chang et al., 2013; J. S. Nomi et al., 2018) and reflect the functional mapping between the subregions and the three intrinsic FNs including the SMN, VAN, and FPN, respectively. The combined approach using topic mapping enabled further characterization of the cognitive and behavioral profiles of the three insular subregions. We found that for both left and right insular, “pain” correlated highly with dAI, PI and vAI, except for left vAI which correlated highly with “working memory” for top term terms. (Figure 4). In a recent meta-analytic study by Ferraro and colleagues, the authors used reverse inference to identify top terms associated with the insular region associated with dysregulations in pain patients and found that “pain or painful” correlated highly with the anterior insular (Ferraro et al., 2021), In accordance with previous studies, the anterior insula is involved in processes related to the acute and chronic pain experience, including empathetic pain ((Fallon, Roberts, & Stancak, 2020; Zhou et al., 2020), and the sensory experience. However, for both dAI and PI our reverse inference emphasized the contribution of the insula to sensory and somatosensory responses. In particular, although specialized pain sensations have been identified for the whole of the insula region, the PI is mostly associated with thermosensory pain (Baier et al., 2014; Craig, Chen, Bandy, & Reiman, 2000). The genetic expressions obtained from the database indicated that specific genetic profiles were predominately expressed in both left and right insula and may suggest their importance in either biological basis for the structural formation or evolutionary functionality. Moreover, although these symbols are well mapped in the database, this current study does not provide the technical meanings related to the structural-functional insular regions. It is, however, important to note that the symbol expressions and their strong correlations are indicative of the fact that while certain gene expressions are strongly associated with specific subregions others are less expressed, such patterns will need to be further investigated as it goes beyond the content of these current study.

By combining both RSFC and DTI and examining their overlaps, we demonstrate a potential approach for investigating the associations between structural and functional connections of any brain system. More importantly, the combined methods enabled the estimation of the relationships between the functional and anatomical connectivity that best describe the insula.

Interestingly, the results echoed that of both functional (Deen et al., 2011) and structural (J. S. Nomi et al., 2018) studies conducted independently. Our study also reflects the importance of the relationship between the two neuroimaging modalities. In addition, meta-analysis expanded and further characterized the insular subregions in terms of both behavioral and cognitive domains as well as their gene expression patterns across specific FNs.

### Limitations

Although the present work represents a comprehensive approach for examining the intersection of RSFC and DTI profiles of the insula, several limitations should be considered. First, issues with model crossing fibers in DTI were inadequately represented, especially within the cerebral cortex, such as in the centrum semiovale (Wedeen et al., 2008). This is a potential problem in Human Connectome Project (HCP) data that could be solved using high-quality imaging, which would enhance fiber tracking in the white matter (Maier-Hein et al., 2019). Secondly, though recent work suggests the existence of gradients of connectivity in the insula (Royer et al., 2020)representing a shift in microstructure, this current work was not able to investigate how this is reflected in the structural-functional correspondence of the insula, further investigation is therefore recommended in this direction.

### Conclusion

Our study combined a data-driven approach and meta-analysis to demonstrate the correspondence between the structural and functional connectivity of insular subregions mapped to well-described intrinsic FNs. Our findings reinforce the tripartite subregions of the insula including the PI, dAI, and vAI, which were significantly mapped to the SMN, VAN, and FPN, respectively. Generally, the significance of our finding firstly answers the question of whether there is a substantial overlap between the DTI and RSFC profiles and if these overlaps are relevant within the scope insula related functions. In this regard, delineating insula subregions that demonstrated a good correspondence between structure and functional profiles provided evidence of the viability of combing the two modalities within the insula system. Secondly, identification of the subregions based on the overlap of the two modalities will provide direct information about the enigmatic subregional properties, thus serving researchers and clinicians to be more specific in their approach in studying the insula and its related disorders. Furthermore, the additional application of the large-scale meta-analysis provided an unbiased approach for characterizing the functional profiles of the subregions and how the anatomical arrangement can further provide better inference for the cognitive and behavioral relations including their gene expression related to the different insular subregions in the human brain.

## Acknowledgments

This work was supported by the National Natural Science Foundation of China (NSFC, No, G0561871420)

## Supplementary Materials and Methods

### Data Acquisition

Details on data acquisition and protocols can be found on the webpage of the human connectome project (https://db.humanconnectome.org). The project was approved by the local Institutional Review Board at Washington University in St. Louis, Missouri USA with informed consent signed by each subject, including the parameters described in (Van Essen et al., 2013). Briefly, the resting-state images were collected using the following parameters: repetition time (TR) = 720 ms; echo time (TE) = 33.1 ms; flip angle = 52°; field of view = 208 ×180 mm^2^; slice number = 72; functional spatial resolution = 2 mm isotropic voxel; multiband factor = 8; echo spacing = 0.58 ms; bandwidth = 2290 Hz/px; and number of volumes (time points) = 1200. whereas the DTI parameters collecting diffusion data consisted a multiband factor of 3, nominal voxel size of 1.25 mm isotropic, 270 diffusion-weighted scans distributed equally over 3 shells (*b* values = 1000, 2000 and 3000 s/mm^2^), and 18 volumes without diffusion weighting (*b* = 0 s/mm^2^).

### Functional data preprocessing

Preprocessing included validated routines, including the removal of the first 10 volumes, head motion-related signal correction with maximum displacements of all subjects being below 3 mm or 3° for translation and rotation, respectively. Each subject’s T1-weighted MPRAGE image was co-registered to the mean functional image with a six-degree-of-freedom linear transformation without re-sampling. Each structural image was segmented into gray matter (GM), white matter (WM), and cerebrospinal fluid (CSF). Linear trends were removed to adjust for signal drift and temporal bandpass filtering (0.01–0.15 Hz) was performed. The functional images were normalized from the individual space to Montreal Neurological Institute (MNI) with voxel size of 2 mm x 2 mm x 2 mm. Several signals of spurious variance were removed by regressing out the following variables: 1) averaged CSF signal of whole-brain; 2) 24 rigid body motion parameters (6 head motion parameters, 6 values at previous time points of 6 head motion parameters, and 12 corresponding squared items); 3) head motion scrubbing regressors identified by framewise displacement (FD) greater than 1 mm (Power, Barnes, Snyder, Schlaggar, & Petersen, 2012), a correction step has been shown to significantly improve the elimination of potential head motion artifacts accompanying signals without altering correlation values (Satterthwaite et al., 2013). We did not smooth nor regress out global mean signal in line with a previous study employing thalamus parcellations (Zhang et al., 2008).

### DTI data preprocessing

We employed a validated preprocessing pipeline (Glasser et al., 2013; P. Wang et al., 2021) which involved: the normalization of the b0 image intensity corrected for EPI distortions, eddy current distortions, and subject movements, and gradient nonlinearities. To facilitate comparison with our RSFC study, preprocessed diffusion data (voxel size: 1.25 ×1.25 ×1.25 m3) were normalized from the individual space to the standard space (MNI space) without changing the spatial resolution using the following procedures: 1) the T1 images were co-registered to the b0 images; 2) the co-registered T1 images were normalized to the MNI space and the transformation matrices were then obtained; 3) diffusion data were warped to the MNI space using these transformation matrices defined above. The parameters of DTI analysis process were used by referring to the operation guide for HCP data in this study (https://www.ismrm.org/15/program_files/Tu03.htm). The whole-brain fiber tracking was performed for each subject using the Fiber Assignment by Continuous Tracking (FACT) algorithm embedded in the Diffusion Toolkit (R. Wang & Benner, 2007). The path tracking was restricted by the predefined turning angle (less than 60°).

## Supplementary Tables

**Table S1.**
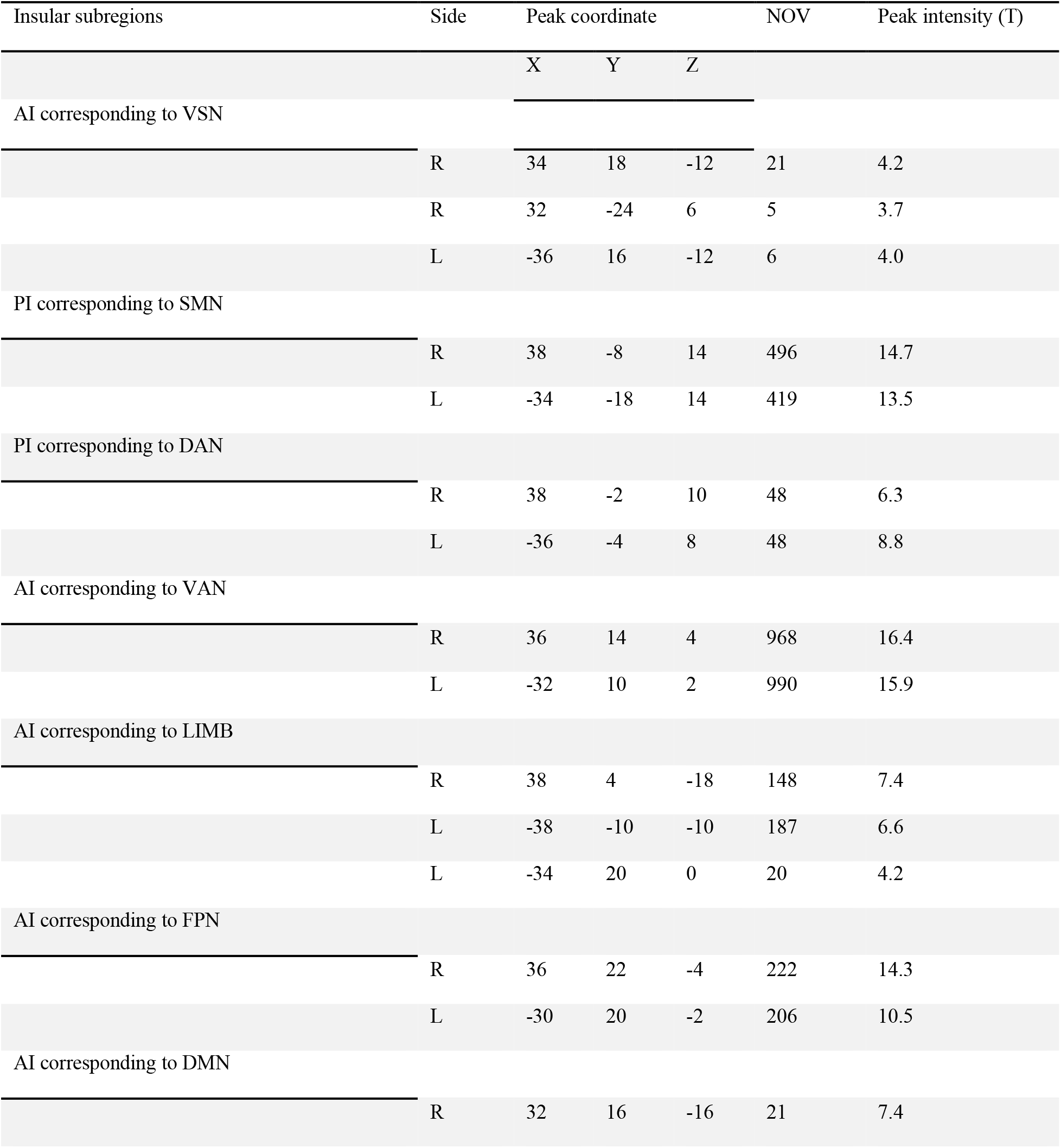
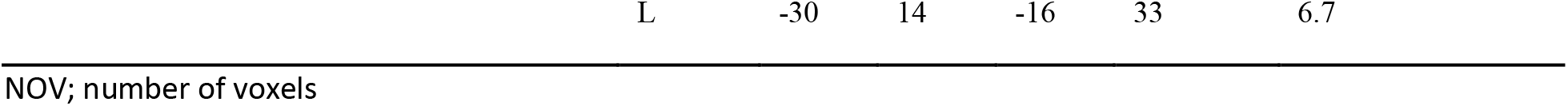
functional insula peak MNI coordinate centers corresponding to each functional network.

**Table S2.** Structural insula peak MNI coordinate centers corresponding to each functional network.

| Insular subregions | Side | Peak coordinate |  |  | NOV | T-value |
| --- | --- | --- | --- | --- | --- | --- |
|  |  | X | Y | Z |  |  |
| AI corresponding to VSN |  |  |  |  |  |  |
|  | R | 33.75 | 9 | -9.5 | 2521 | 65 |
|  | L | -35 | -11 | -8.25 | 3658 | 77 |
| PI corresponding to SMN |  |  |  |  |  |  |
|  | R | 33.75 | -12.25 | 11.75 | 3528 | 85 |
|  | L | -35 | -14.75 | -2 | 3807 | 102 |
| AI corresponding to DAN |  |  |  |  |  |  |
|  | R | 36.25 | 6.5 | -5.75 | 3003 | 53 |
|  | L | -35 | 1.5 | -5.75 | 3256 | 66 |
| AI corresponding to VAN |  |  |  |  |  |  |
|  | R | 31.25 | 15.25 | -3.25 | 3620 | 97 |
|  | L | -35 | 1.5 | -7 | 3954 | 109 |
| AI corresponding to LIMB |  |  |  |  |  |  |
|  | R | 31.25 | 7.75 | -<br>13.25 | 1612 | 92 |
|  | L | -31.25 | 7.75 | -12 | 2885 | 101 |
| AI corresponding to FPN |  |  |  |  |  |  |
|  | R | 31.25 | 15.25 | -3.25 | 3194 | 97 |
|  | L | -33.75 | 10.25 | -7 | 3587 | 97 |
| AI corresponding to DMN |  |  |  |  |  |  |
|  | R | 33.75 | 11.5 | -7 | 3256 | 93 |
|  | L | -33.75 | 10.25 | -7 | 3746 | 98 |

**Table S3.**
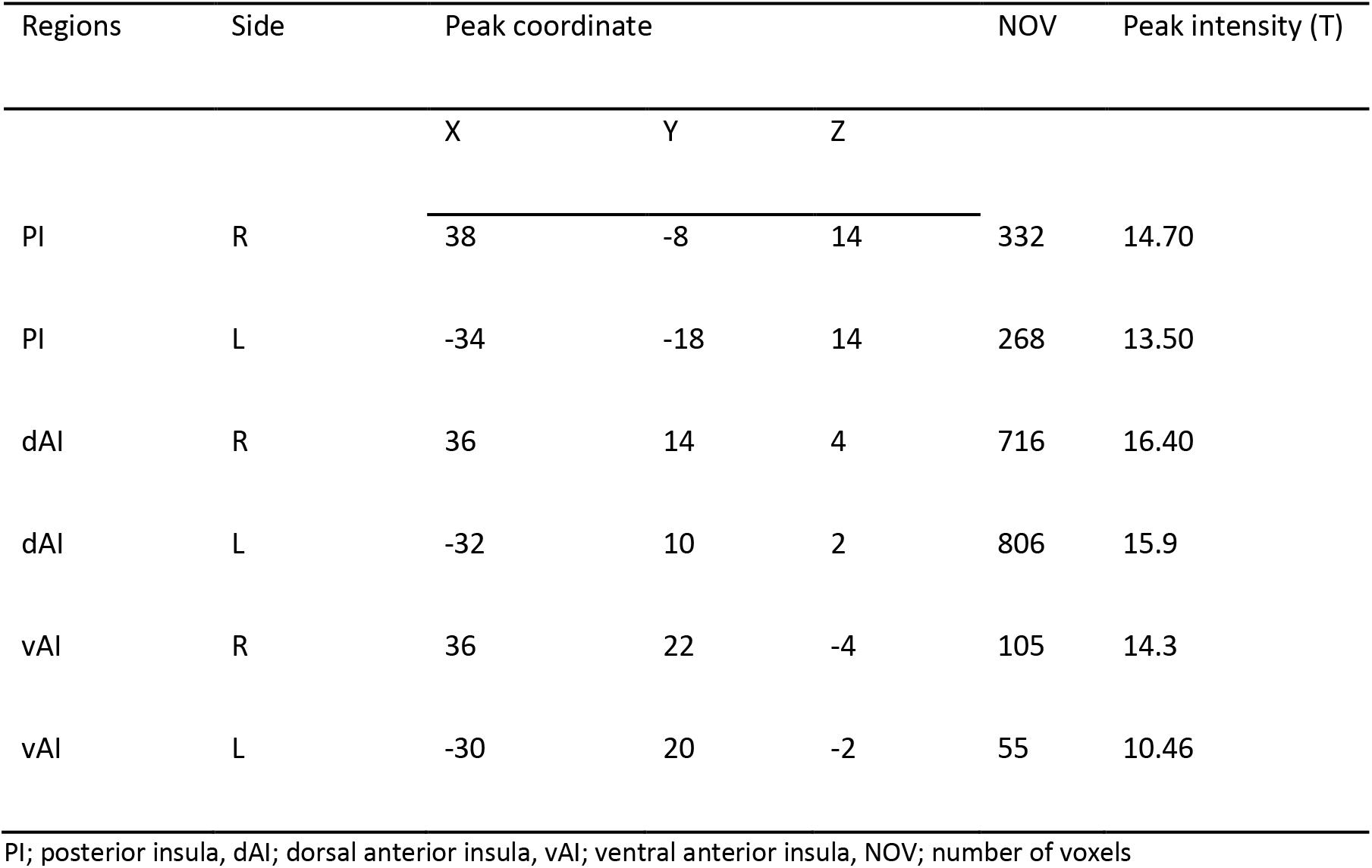
Peak coordinates of insular subregions in MNI space

| Regions | Side | Peak coordinate |  |  | NOV | Peak intensity (T) |
| --- | --- | --- | --- | --- | --- | --- |
|  |  | X | Y | Z |  |  |
| PI | R | 38 | -8 | 14 | 332 | 14.70 |
| PI | L | -34 | -18 | 14 | 268 | 13.50 |
| dAI | R | 36 | 14 | 4 | 716 | 16.40 |
| dAI | L | -32 | 10 | 2 | 806 | 15.9 |
| vAI | R | 36 | 22 | -4 | 105 | 14.3 |
| vAI | L | -30 | 20 | -2 | 55 | 10.46 |

PI; posterior insula, dAI; dorsal anterior insula, vAI; ventral anterior insula, NOV; number of voxels

